# Intrinsic Timescales Across the Basal Ganglia

**DOI:** 10.1101/2021.04.07.438040

**Authors:** Simon Nougaret, Valeria Fascianelli, Sabrina Ravel, Aldo Genovesio

**Author notes:** **Contact Info** Corresponding authors are noted with an asterisk in the author list. Correspondence: Simon Nougaret, Aldo Genovesio. **Email address of each author,** Simon Nougaret, Valeria Fascianelli, Sabrina Ravel, Aldo Genovesio.

## Abstract

Recent studies have shown that neuronal stability over time can be estimated by the structure of the spike-count autocorrelation of neuronal populations. This estimation, called the intrinsic timescale, has been computed for several cortical areas and can be used to propose a cortical hierarchy reflecting a scale of temporal receptive windows between areas. In this study, we performed an autocorrelation analysis on neuronal populations of three basal ganglia (BG) nuclei, including the striatum and the subthalamic nucleus (STN), the input structures of the BG, and the external globus pallidus (GPe). The analysis was performed during the baseline period of a motivational visuomotor task in which monkeys had to apply different amounts of force to receive a different amount of reward. We found that the striatum and the STN have longer intrinsic timescales than the GPe. Moreover, our results allow for the placement of these subcortical structures within the already-defined scale of cortical temporal receptive windows. Estimates of intrinsic timescales are important in adding further constraints in the development of computational models of the complex dynamics among these nuclei and throughout cortico-BG-thalamo-cortical loops.

## Introduction

Organization of the brain has been described following different principles. For example, areas can be organized based on the laminar pattern of origins and terminations of cortico-cortical projections (Barbas and Rempel-Clower, 1997; Felleman and Van Essen, 1999) or based on topological projection sequences (Petroni et al., 2001). Following a proposed anatomical hierarchy of the visual, somatosensory, and motor cortices (Felleman and Van Essen, 1999), and considering the laminar structure of the prefrontal cortico-cortical projections, prefrontal areas are at the top of this hierarchy (Murray et al., 2014). Interestingly, this anatomical hierarchy is mirrored by the intrinsic fluctuations in spiking activity across these areas at rest (Murray et al., 2014; Ogawa and Komatsu, 2010). Computed from their spike-count autocorrelation, these intrinsic timescales are considered to be a measure of neuronal stability. By comparing different cortical areas, past studies (Cirillo, Fascianelli et al., 2018; Ogawa and Komatsu, 2010) have shown that prefrontal areas have the longest timescales, the posterior parietal and the dorsal premotor cortex have intermediate timescales, and the somatosensory cortex has the shortest timescale. The proposed cortical hierarchy (Chen et al., 2015; Ogawa and Komatsu, 2010) is intended to reflect a scale detailing temporal receptive windows with higher-level areas with the longest timescales representing the progressive accumulation of neuronal inputs and supporting high-level cognitive decision-making processes.

All cortical areas except for the primary visual and auditory areas project to the basal ganglia (BG), which serve as the substrate of several cognitive processes such as context- and value-based decision-making, reinforcement learning, inhibition control, and working memory (Mink, 1996). Here, we analyzed the intrinsic timescales of neuronal populations in the striatum (phasically active neurons, PANs, or putative projection neurons), the subthalamic nucleus (STN), and the external globus pallidus (GPe) of macaque monkeys during the baseline period of a visuomotor task (Nougaret and Ravel, 2015, 2018). We found that the input structures of the BG, the striatum and the STN, exhibited longer timescales than the GPe. Describing the differences between the timescales of these populations can help lead to a better understanding of the functional specialization of these structures and validate computational models of action selection.

## Results

We analyzed neuronal activity during the baseline period of a visuomotor task as described by Nougaret and Ravel (2015, 2018). During the 1 s baseline period, monkeys maintained a basal pressing force on a lever while waiting for the presentation of a pair of visual stimuli that informed them of the amount of force required and the amount of reward they would receive upon completion of the trial. We computed the spike-count autocorrelation structure for each neuron as a function of time lag, and estimated its decay constant (intrinsic timescale τ) with an exponential fit. We assigned the intrinsic timescale to the whole neuronal population and to single neurons as described in Materials and Methods. The database we analyzed consisted of 78 neurons recorded in the STN (30 and 48 from monkey M and monkey Y, respectively); 158 PANs (96 and 62 from monkey M and monkey Y, respectively) recorded in the striatum, presumed to be medium spiny projection neurons (Inokawa et al., 2010); and 92 irregular neurons, corresponding to high-frequency discharge neurons (HFD; DeLong, 1971), from the GPe (41 and 51 from monkey M and monkey Y, respectively). Only neurons from the GPe were analyzed in a previous study (Nougaret and Ravel, 2018). Localizations of PANs in the striatum and STN neurons were assessed as in previous studies (Nougaret and Ravel, 2015, 2018) using MRI scans with electrodes for locating trajectories, from which the neurons were recorded.

### Intrinsic timescales of STN neurons, PANs, and GPe neurons

To assess the spike-count autocorrelation values as a function of time lags, a non-zero mean activity for each neuron in each 50 ms bin during the baseline period was required. In particular, 77/78 neurons in the STN, 103/158 PANs in the striatum, and 92/92 neurons in the GPe fulfilled this requirement (see Materials and Methods). Figure 1 (left) shows the autocorrelation values as a function of time lags averaged across neurons for each brain structure, with the exponential fit superimposed along with the estimated timescale τ. In particular, the GPe showed a shorter timescale (τ_GPe ± sem_GPe = (120 ± 3) ms, Figure 1C) than both the striatum (τ_PANs ± sem_PANs = (258 ± 35) ms; Figure 1B) within the error (τ_PANs − τ_GPe ± Δ(τ_PANs − τ_GPe) = (138 ± 35) ms) and the STN (τ_STN ± sem_STN = (230 ± 86) ms; Figure 1A) within the error (τ_STN − τ_GPe ± Δ(τ_STN − τ_GPe) = (110 ± 86) ms). Moreover, the intrinsic timescale of the STN is compatible with the timescale of the striatum within the error (τ_PANs − τ_STN ± Δ (τ_PANs − τ_STN) = (28 ± 93) ms).

**Figure 1:**
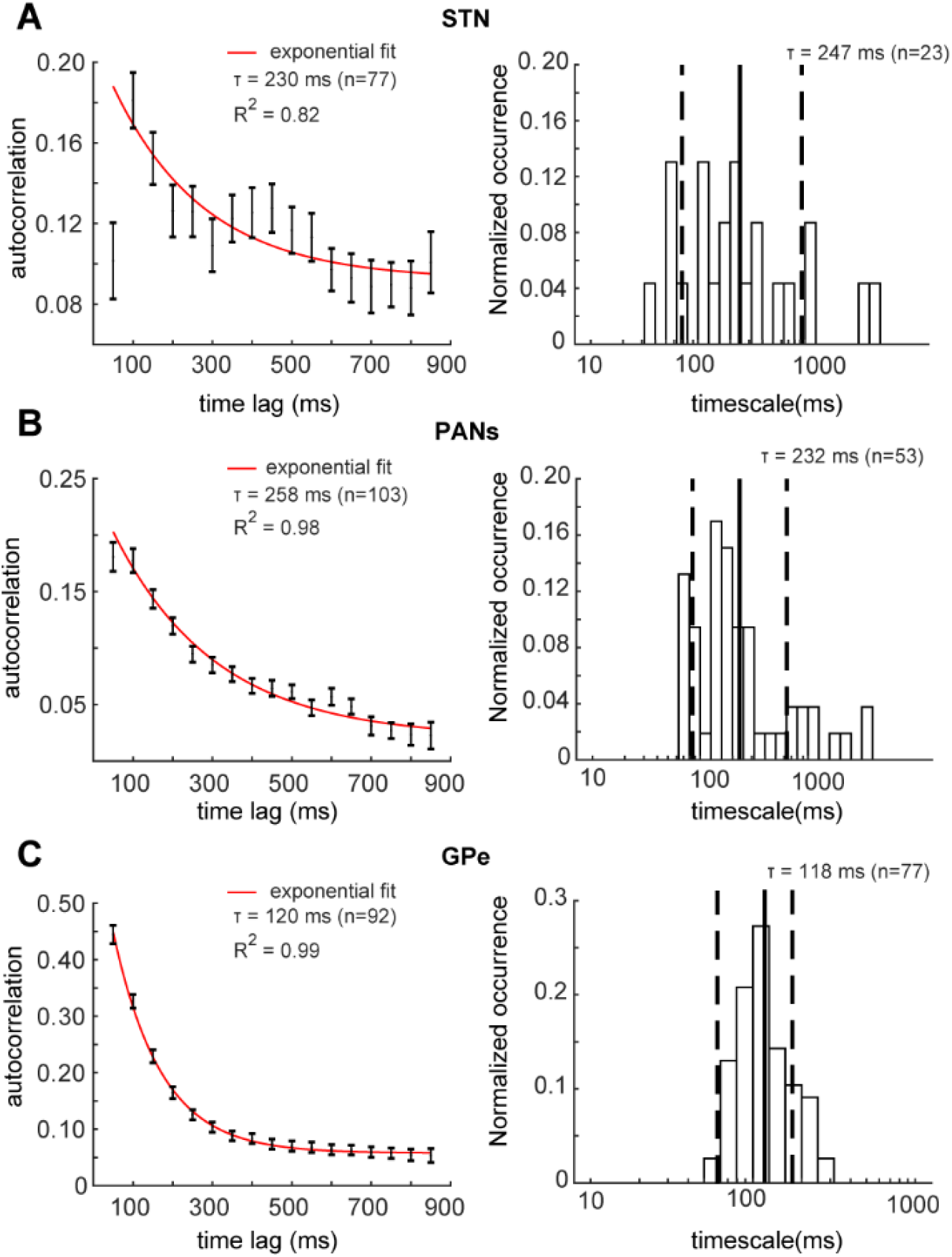
Mean autocorrelation values and single timescale distribution. **A)** Left Panel: mean autocorrelation averaged across all neurons (n = 77) recorded in the subthalamic nucleus (STN) using 50 ms time bins in a 900 ms time window during the baseline period (mean ± SEM). The solid red line is the exponential fit. The autocorrelation at 50 ms has been excluded from the fit procedure. The intrinsic timescale τ is shown in the top right corner, with the R^2^ value as a goodness of fit estimator. Right panel: single neuron timescale distribution (n = 23) computed in the same baseline period as in the population timescale shown on the left panel. The solid and dashed lines are the mean(log(τ)) ± SD (log(τ)). The mean of the timescale distribution is shown in the top right corner. **B)** Left Panel: mean autocorrelation averaged across neurons (n = 103) recorded in phasically active neurons (PANs) of the striatum in the same baseline period as A). The autocorrelation value at 50 ms has been excluded from the fit procedure as in A). Right panel: single neuron timescale distribution (n = 53). **C)** Left Panel: mean autocorrelation averaged across neurons (n = 92) recorded in the external globus pallidus (GPe) in the same baseline period as A) and B). The autocorrelation values at all time lags have been included in the fit procedure. Right panel: single neuron timescale distribution (n = 77).

Comparing the timescales of the BG structures, we found that the estimated GPe timescale was more accurate than the timescale estimates for the other two BG structures. This result could be explained by less heterogeneity in the autocorrelation structures of single neurons compared to the other two BG structures. We then wanted to confirm the results found at the population level at the single-cell level. For this, we computed the timescale for each neuron (Figure 1, right) satisfying the requirements detailed in (1) and (4) in Materials and Methods. Following this selection process, we kept 23/78 neurons from the STN, 53/158 neurons from the striatum, and 77/92 neurons from the GPe. Even when selecting a subset of neurons, the results at the single-cell level were similar to the previous population analysis. The distribution of the timescale values of GPe neurons was significantly lower than the timescale distributions of STN neurons (Mann-Whitney test, p = 0.0074) and PANs (Mann-Whitney test, p = 4 × 10^−5^). No significant difference was found between STN and PAN timescale distributions (Mann-Whitney test, p = 0.9369).

To investigate the degree of heterogeneity in the single timescale distributions for each BG structure, we computed the coefficient of variation (CV) of the timescale distribution using formula (5) in Materials and Methods. We found a higher degree of heterogeneity in STN neuron (CV = 23%) and PAN (CV = 19%) τ distributions than in the GPe neuron τ distribution (CV = 8%), in line with previous population analysis results.

## Discussion

To our knowledge, our study is the first to report estimations of intrinsic timescales of neuronal populations at the subcortical level, and reveals timescale differences between the input structures of the BG, the striatum and the STN, and the GPe. Earlier studies have used the autocorrelation function to understand the firing pattern properties of single cells within the BG (Bar-Gad et al., 2002; Magill et al., 2000, 2001) and midbrain dopaminergic neurons (Paladini and Tepper, 2016). Specifically, autocorrelograms of single cells have been used to classify these cells into different subpopulations, for example into GPe neurons (Bugaysen et al., 2010) or different types of dopaminergic neurons (Paladini and Tepper, 2016), and to assess firing rate rhythmicity of single neurons from BG nuclei in healthy and diseased conditions (Heimer et al., 2002; Magill et al., 2000, 2001; Raz et al., 2000). In this study, we used the autocorrelation function to characterize the properties of neuronal populations in the BG and place them in the context of already-known intrinsic timescales throughout the cortex.

We found that the striatum and the STN exhibited longer timescales (258 and 230 ms respectively) compared to the GPe (120 ms). A cortical hierarchy has already been described (Murray et al., 2014) based on values from seven cortical areas (Figure 2, light gray circles), placing the prefrontal areas, anterior cingulate cortex (ACC; average value = 303 ms), orbitofrontal cortex (OFC; average value = 182 ms), and lateral prefrontal cortex (LPFC; average value = 166 ms) at the top of this hierarchy with the longest timescale values. The same study then reported intermediate timescales in the lateral intraparietal cortex (LIP; average value = 114.5 ms) and the secondary somatosensory cortex (S2), and the shortest timescales in the medio-temporal area (MT) of the visual cortex and the primary somatosensory cortex (S1). Other studies (Cavanagh et al., 2016, medium gray circles; Fascianelli et al., 2019, dark gray circles) later confirmed the previous results overall, reporting comparable LPFC (231/248 ms), OFC (241/190 ms), and ACC (332 ms) timescale values. Genovesio and colleagues (Cirillo, Fascianelli et al., 2018; Fascianelli et al., 2019) then extended the hierarchy previously described by assigning intrinsic timescales to the frontopolar cortex (PFp; 242 ms) and the dorsal premotor cortex (PMd; 131 ms).

**Figure 2:**
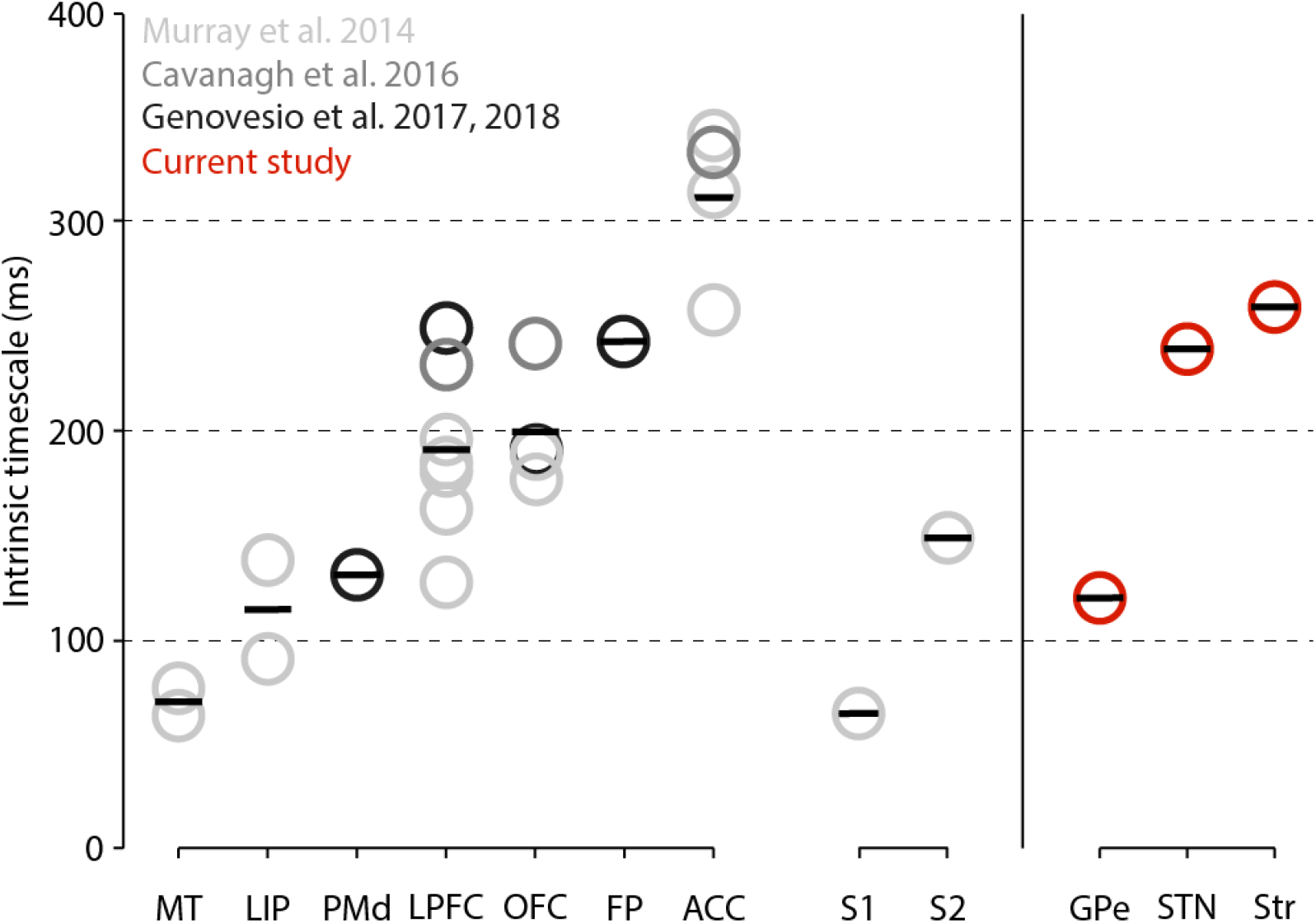
Hierarchical organization of intrinsic timescales of cortical and subcortical structures. Left Panel: Intrinsic timescales of nine cortical areas reported by Murray et al. (2014) in light gray, by Cavanagh et al. (2016) in medium gray, and by Genovesio and collegues (Cirillo, Fascianelli, 2018; Fascianelli et al., 2019) in dark gray. The seven areas on the left (MT, LIP, PMd, LPFC, OFC, FP, and ACC) are part of the visual and prefrontal cortices. The two areas on the right are part of the somatosensory cortex (S1, S2). Each circle represents the average τ for each cortical area reported in each study. Each bar represents the average τ among the studies. Right Panel: Same representation for the three subcortical structures (GPe, STN, and striatum) analyzed in the present study. Abbreviations: ACC, anterior cingulate cortex; FP, frontopolar cortex; GPe, external globus pallidus; LIP, lateral intraparietal cortex; LPFC, lateral prefrontal cortex; MT, medio-temporal area (of visual cortex); OFC, orbitofrontal cortex; PMd, dorsal premotor cortex; S1, primary somatosensory cortex; S2, secondary somatosensory cortex; STN, subthalamic nucleus.

The striatum and STN intrinsic timescales reported here place these structures on a comparable level as the timescales assigned to prefrontal areas. Importantly, in both of these BG structures recordings were mainly from their associative parts, known for receiving their major inputs from the prefrontal cortex (Alexander et al., 1986; Haber and Knutson, 2010; Haynes and Haber, 2013; Nougaret et al., 2013). In contrast, the GPe exhibited an intrinsic timescale of only 120 ms, which places it at the same intermediate level as the PMd, LIP, and S2. One possibility is that this lower timescale could reflect lower temporal information integration ability of the GPe among BG structures. Our study shows a gradient of receptive time windows within BG circuitry (Figure 2, red circles). As suggested for areas of the visual system (Ogawa and Komatsu, 2010), the differences found among the BG nuclei could reflect a broader window of temporal information storage in the striatum and the STN compared to in the GPe. Indeed, intrinsic timescales could be described as “a temporal counterpart of the spatial hierarchy” (Chen et al., 2015). Previous studies (Chaudhuri et al., 2015; Murray et al., 2014) shed light on the existence of a parallel between the anatomical hierarchy of cortical areas and their intrinsic timescales. Drawing a parallel with the cortex, the interpretation of a functional hierarchy based on timescales described for cortical areas might apply to the BG. This would suggest a greater need for information accumulation within the BG input nuclei rather than in the GPe. The convergence of information from multiple cortical areas could explain the necessity for longer timescales in both input structures because of the need of striatal projection neurons and subthalamic neurons to gate and maintain information from prefrontal neurons (Frank et al., 2006; O’Reilly and Frank, 2006).

A large body of work in primate neurophysiology has shown that, at the neural level, choosing corresponds to crossing a firing rate threshold in the cortex. Depending on the environment and on the decision being made, the threshold level can be regulated. A computational study (Lo and Wang, 2006) implemented a biophysically-based network model of decision thresholds of the cortical-BG-superior colliculus (SC) pathway. After comparing the different nodes of these networks, the authors concluded that the all-or-none activity of SC neurons is triggered by a threshold crossed by cortical neurons that can be optimally tuned by the strength of cortico-striatal synapses. Indeed, through this pathway, the output structures of the BG inhibit cortex and SC activity to preclude inappropriate motor outputs (Jahanshahi et al., 2015). Ding and Gold (2013) have hypothesized that “the BG may convert cortical representations of sensory evidence into evaluative quantities”, allowing generation and adjustment of decisions. They suggest that the BG can modify the decision rules by modifying the decision threshold, but also modify the “value of a developing decision variable”. Different BG models hypothesize that the main role of the BG nuclei is to act as a central selection device (Redgrave et al., 1999) that examines each action requested based on its urgency and salience (Bogacz and Gurney, 2007) and that, with their unique anatomical properties, allows the allocation of motor resources to the appropriate actions. The striatum and the STN have distinct roles in these processes. According to Frank and colleagues, the former has a crucial role in gating sensory input for updating working memory in the prefrontal cortex and then in maintaining it, preventing the influence of distracting information (O’Reilly and Frank, 2006), especially when adaptive gating is necessary for the processing of multiple goal demands. The STN is supposed to act as a brake, particularly during high-conflict decisions, reducing premature responses and refining the selection process that takes place via cortico-striatal pathways (Cavanagh et al., 2011; Frank, 2006, 2007). Both functions reflect the need to accumulate information over time, and support the long intrinsic timescales exhibited by the input structures of the BG.

On the other hand, models of BG action selection assign another role to GPe neurons. Gurney and colleagues (Bogacz and Gurney, 2007; Gurney et al., 2001) have proposed that the GPe, mainly through its massive projections into the STN, automatically limits the activity of BG output structures and allow the network to make a selection. The control that the GPe exerts on other BG nuclei is supported by *in vitro* electrophysiological studies showing that the firing rate of GPe neurons can be approximated by a linear function of the injected current (Namu and Llinaś, 1994). In contrast, the firing of the STN neurons fit better with an exponential function of their inputs (Bogacz and Gurney, 2007). This understanding is also consistent with *in vivo* electrophysiological studies in primates that report independent encoding of task variables by GPe neurons (Arkadir et al. 2004; Nougaret and Ravel, 2018), suggesting more of a parallel processing of information by GPe neurons rather than an integration of different variables (Nougaret and Ravel, 2018). Taken together, these results are in line with the intermediate intrinsic timescale exhibited by the GPe in the present study. However, a recent computational study (Chaudhuri et al., 2015) suggests caution around assigning timescales to brain areas too rigidly, because processing different sensory inputs may lead to different timescales based on their model. Moreover, the three BG nuclei studied here exhibited different degrees of heterogeneity in their single unit timescales (Figure 1, left panel). In particular, the input structures displayed a higher degree of heterogeneity than the GPe. Some studies have shown that within each cortical area, the individual intrinsic timescale computed during a baseline period predicted the strength of response modulation during following task periods in the LIP (Nishida et al., 2014), PMd (Cirillo, Fascianelli et al., 2018), and dlPFC (Fascianelli et al., 2019), suggesting that neurons with longer timescales are more involved in the encoding of task-related information. We could hypothesize that the high degree of heterogeneity found in the input structures of the BG could serve to support the heterogeneity of information that these structures have to process with different temporal integration requirements, although further study is necessary to reach conclusions about these functions.

Our study is the first to quantify intrinsic timescales of BG nuclei, and some limitations should be considered. First, our datasets are relatively small compared to others used for the cortex and have high variability at the population level. This is mainly true for the input structures, which showed a higher degree of heterogeneity at the single-cell timescale level. Second, the BG nuclei are known to be partially specialized in sensorimotor, associative, and limbic territories, and our datasets cover mainly the associative and the limbic parts of these structures. For future studies, it remains to be investigated how our BG nuclei timescale estimates could be generalized at the whole-structure level, and whether our estimates are consistent with the timescales of other BG nuclei not studied here. It is also important to investigate the relationships within each structure at the single-cell level between timescales and show persistent representations of task-relevant signals, as has been done in the cortex (Cavanagh et al., 2016; Cirillo, Fascianelli et al., 2018; Fascianelli et al., 2019; Nishida et al., 2014). We believe that notwithstanding these limitations, the timescales reported here could be useful as a first approximation for the validation of computational models of action selection based on evidence accumulated through cortico-striatal/subthalamic synapses and architecture of cortico-BG-cortical loops, as has been done for the cortex (Chaudhuri et al., 2015).

## Author Contributions

S.R. designed the experiment, S.N. and S.R collected the data, V.F. performed the timescales analysis, S.N, S.R., V.F. and A.G. interpreted the results and wrote the manuscript, S.R. supervised the experimental part of the study, A.G. supervised the analysis part of the study.

## Acknowledgment

This work was supported by Centre National de la Recherche Scientifique, the Aix-Marseille Université, the Fondation de France (Parkinson’s Disease Program Grant 2008 005902 to SR), the project H2020 (Grant PH1181642DB714F6 to AG).

## Materials and Methods

### Resource availability

Further information and requests for resources and reagents should be directed and will be fulfilled by Simon Nougaret.

The datasets and code supporting the current study have not been deposited in a public repository yet, but are available from the corresponding author on request.

### Experimental model and subject details

Two male rhesus monkeys (*Macaca mulatta*) were used in this study. Relevant details about the animals have already been reported in Nougaret and Ravel (2015, 2018).

### Method details

#### Dataset

The experimental procedures followed French laws on animal experimentation, the European directive on animal protection, and the National Institute of Health’s Guide for the Care and Use of Laboratory Animals. The experimental details of the datasets used in the current study have already been reported (Nougaret and Ravel, 2015, 2018). The single-unit activity of two male rhesus monkeys (*Macaca mulatta*), recorded from three populations of neurons within the (BG), were analyzed during a foreperiod, considered as a baseline period in which no cognitive process was engaged. During this 1 s period, both monkeys needed to maintain a basal pressing force on a lever and wait for the presentation of a pair of visual stimuli indicating the amount of force needed and the amount of reward to be expected at the end of the trial. The data set consisted of 1) a population of 158 putative projection neurons recorded in the striatum, also called phasically active neurons (PANs); 2) a population of 78 neurons from the subthalamic nucleus (STN); and 3) a population of 92 irregular neurons from the external globus pallidus (GPe).

### Quantification and statistical analysis

#### Spike-count autocorrelation structure

We analyzed the recorded activity of neurons in the STN, GPe, and striatum (PANs) from 100 ms after the beginning of the trial until the end of the baseline period for a total of 900 ms of baseline activity. The same baseline period had already been chosen (Nougaret and Ravel, 2015) for electrophysiological analysis of neurons recorded for the same task. We included only correct trials in the following analyses. For each structure, we selected neurons satisfying the following criterium:

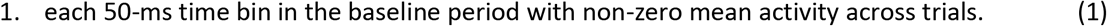

We performed all analyses with MatLab (The MathWorks, Inc., Natick, MA, USA).

To assess the spike-count autocorrelation structure, we calculated the spike count during the baseline period in 50 ms time bins. It is worth noting that the results did not change within a difference of 20% of the bin length. Given a neuron, the spike-count autocorrelation value across trials between time bins *k* and *j* (*k, j* as integer numbers) at a time lag equal to |*k-j*|x *Δ* (*Δ* = 50 ms), the Pearson’s correlation coefficient *r* is defined as follows (Murray et al., 2014):

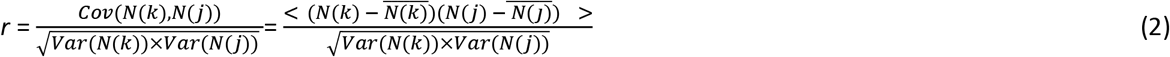

where *N(k)* and *N(j)* are the spike counts computed in the *k* and *j* time bins, respectively, and 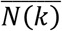 and 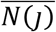 are the spike counts averaged across trials in *k* and *j* time bins, respectively. The covariance (*Cov*), the variance (*Var*), and the autocorrelation value *r* were computed for each possible combination of pair-bins (*k, j*). We calculated the autocorrelation values as a function of the time lags for each neuron satisfying the criterion in (1). We subsequently computed the autocorrelation structure for the whole neuronal population by averaging the coefficient *r* across neurons at a fixed time lag. We obtained the autocorrelation values as a function of time lags for the entire population, and we performed an exponential fit as defined below (Murray et al., 2014):

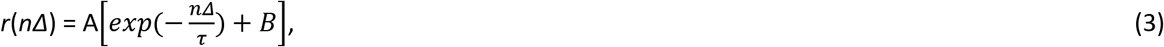

where *nΔ* indicates the time lag between the time bins *k* and *j*, with *n* = |*k* - *j* | (*n*= 1,2, …, 18); *r* is the autocorrelation value at time lag *nΔ*; A is the amplitude; τ is the decay constant of the exponential function, called intrinsic timescale; and B is the offset that mirrors the value of *r* in the limit of time lag *nΔ*→∞ (i.e., time lag values much larger than our 900 ms baseline length). Throughout this paper, we refer to intrinsic timescale as simply timescale or τ.

#### Single-neuron intrinsic timescale

We also fit each single spike-count autocorrelation decay with the exponential function in (3) to estimate the single-neuron intrinsic timescale. We further reduced the neuronal sample by selecting those neurons satisfying both the criterion in (1) and the following requirements:

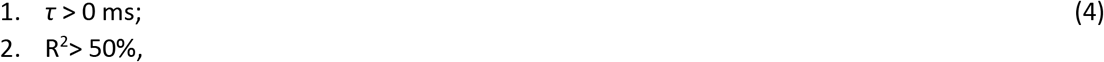

where R^2^ is the coefficient of determination obtained by the fit. The first requirement was introduced because a negative or 0 ms *τ* value is meaningless; the second requirement of an R^2^ larger than 50% was a trade-off between the need to keep as many neurons as possible and the importance of having a good fit. We also excluded outliers, defined as neurons having an intrinsic timescale below the 5th percentile and above the 95th percentile of the *τ* distribution, from the sample. This last requirement was established due to the heterogeneity of timescale values within each brain structure and to avoid having an estimate of the mean of the intrinsic timescales biased towards the outlier values. We further investigated the degree of heterogeneity of the single timescale distribution for each brain structure. To quantify this, we used the coefficient of variation (CV), defined as follows:

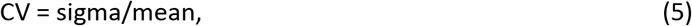

where sigma and mean are the standard deviation and mean of the timescale distribution, respectively.

## References

Alexander, G., DeLong, M.R., and Strick, P. (1986). Parallel Organization of Functionally Segregated Circuits Linking Basal Ganglia and Cortex. Annual Review of Neuroscience 9, 357–381. https://doi.org/10.1146/annurev.ne.09.030186.002041

Arkadir, D., Morris, G., Vaadia, E., and Bergman, H. (2004). Independent coding of movement direction and reward prediction by single pallidal neurons. Journal of Neuroscience 24, 10047–10056. DOI: 10.1523/JNEUROSCI.2583-04.2004

Barbas, H., and Rempel-Clower, N. (1997). Cortical structure predicts the pattern of corticocortical connections. Cerebral Cortex 7, 635–646. https://doi.org/10.1093/cercor/7.7.635

Bar-Gad, I., Ritov, Y., and Bergman, H. (2002). The High Frequency Discharge of Pallidal Neurons Disrupts the Interpretation of Pallidal Correlation Functions. In: Nicholson, L.F.B., Faull, R.L.M. (eds) The Basal Ganglia VII. Advances in Behavioral Biology, vol 52, 35–42. Springer, Boston, MA. https://doi.org/10.1007/978-1-4615-0715-45

Bogacz, R., and Gurney, K. (2007). The basal ganglia and cortex implement optimal decision making between alternative actions. Neural Computation 19, 442–477. DOI: 10.1162/neco.2007.19.2.442

Bugaysen, J., Bronfeld, M., Tischler, H., Bar-Gad, I., and Korngreen, A. (2010). Electrophysiological characteristics of globus pallidus neurons. PLoS ONE 5:e12001. https://doi.org/10.1371/journal.pone.0012001

Cavanagh, J.F., Wiecki, T. V., Cohen, M.X., Figueroa, C.M., Samanta, J., Sherman, S.J., and Frank, M.J. (2011). Subthalamic nucleus stimulation reverses mediofrontal influence over decision threshold. Nature Neuroscience 14, 1462–1467. DOI: 10.1038/nn.2925

Cavanagh, S.E., Wallis, J.D., Kennerley, S.W., and Hunt, L.T. (2016). Autocorrelation structure at rest predicts value correlates of single neurons during reward-guided choice. eLife 5, e18937. https://doi.org/10.7554/eLife.18937.001

Cirillo, R., Fascianelli, V., Ferrucci, L., and Genovesio, A. (2018). Neural Intrinsic Timescales in the Macaque Dorsal Premotor Cortex Predict the Strength of Spatial Response Coding. iScience 10, 203–210. https://doi.org/10.1016/j.isci.2018.11.033.

Chaudhuri, R., Knoblauch, K., Gariel, M.A., Kennedy, H., and Wang, X.J. (2015). A Large-Scale Circuit Mechanism for Hierarchical Dynamical Processing in the Primate Cortex. Neuron 88, 419–431. http://dx.doi.org/10.1016/j.neuron.2015.09.008

Chen, J., Hasson, U., and Honey, C.J. (2015). Processing Timescales as an Organizing Principle for Primate Cortex. Neuron 88, 244–246. http://dx.doi.org/10.1016/j.neuron.2015.10.010.

DeLong, M.R. (1971). Activity of Pallidal Neurons During Movement. Journal of Physiology 34, 414–27. DOI: 10.1152/jn.1971.34.3.414

Ding, L., and Gold, J.I. (2013). The basal ganglia’s contributions to perceptual decision making. Neuron 79, 640–649. http://dx.doi.org/10.1016/j.neuron.2013.07.042

Fascianelli, V., Tsujimoto, S., Marcos, E., and Genovesio, A. (2019). Autocorrelation Structure in the Macaque Dorsolateral, but not Orbital or Polar, Prefrontal Cortex Predicts Response-Coding Strength in a Visually Cued Strategy Task. Cerebral Cortex 29, 1–12. DOI: 10.1093/cercor/bhx321

Felleman, D.J., and Van Essen, D.C. (1991). Distributed hierarchical processing in the primate cerebral cortex. Cerebral Cortex 1, 1–47. https://doi.org/10.1093/cercor/1.1.1

Frank, M.J. (2006). Hold your horses: a dynamic computational role for the subthalamic nucleus in decision making. Neural networks : the official journal of the International Neural Network Society 19, 1120–36. DOI: 10.1016/j.neunet.2006.03.006

Frank, M.J. (2007). Hold Your Horses: Impulsivity, Deep Brain Stimulation, and Medication in Parkinsonism. Science 1309, 1309–1312. http://www.ncbi.nlm.nih.gov/pubmed/17962524.

Gurney, K., Prescott, T.J., and Redgrave, P. (2001). A computational model of action selection in the basal ganglia. I. A new functional anatomy. Biological Cybernetics 84, 401–410. DOI: 10.1007/PL00007984

Haber, S.N., and Knutson, B. (2010). The reward circuit: Linking primate anatomy and human imaging. Neuropsychopharmacology 35, 4–26. https://doi.org/10.1038/npp.2009.129

Haynes, W.I. a, and Haber, S.N. (2013). The organization of prefrontal-subthalamic inputs in primates provides an anatomical substrate for both functional specificity and integration: implications for Basal Ganglia models and deep brain stimulation. Journal of Neuroscience 33, 4804–14. DOI: 10.1523/JNEUROSCI.4674-12.2013

Heimer, G., Bar-Gad, I., Goldberg, J.A., and Bergman, H. (2002). Synchronization of Pallidal Activity in The Mptp Primate Model of Parkinsonism is not Limited to Oscillatory Activity. In: Nicholson, L.F.B., Faull, R.L.M. (eds) The Basal Ganglia VII. Advances in Behavioral Biology, vol 52, 29–34. Springer, Boston, MA. https://doi.org/10.1007/978-1-4615-0715-44

Inokawa, H., Yamada, H., Matsumoto, N., Muranishi, M., and Kimura, M. (2010). Juxtacellular labeling of tonically active neurons and phasically active neurons in the rat striatum. Neuroscience 168, 395–404. DOI: 10.1016/j.neuroscience.2010.03.062

Jahanshahi, M., Obeso, I., Rothwell, J.C., and Obeso, J.A. (2015). A fronto-striato-subthalamic-pallidal network for goal-directed and habitual inhibition. Nature Reviews Neuroscience 16, 719–732. https://doi.org/10.1038/nrn4038

Lo, C.C., and Wang, X.J. (2006). Cortico-basal ganglia circuit mechanism for a decision threshold in reaction time tasks. Nature Neuroscience 9, 956–963. https://doi.org/10.1038/nn1722

Magill, P.J., Bolam, J.P., and Bevan, M.D. (2000). Relationship of Activity in the Subthalamic Nucleus– Globus Pallidus Network to Cortical Electroencephalogram. Journal of Neuroscience 20, 820–833. DOI: 10.1523/JNEUROSCI.20-02-00820.2000

Magill, P.J., Bolam, J.P., and Bevan, M.D. (2001). Dopamine regulates the impact of the cerebral cortex on the subthalamic nucleus–globus pallidus network. Neuroscience 20, 820–833. DOI: 10.1016/s0306-4522(01)00281-0

Mink, J.W. (1996). The basal ganglia: focused selection and inhibition of competing. Progress in Neurobiology 50, 381–425. DOI: 10.1016/s0301-0082(96)00042-1

Murray, J.D., Bernacchia, A., Freedman, D.J., Romo, R., Wallis, J.D., Cai, X., Padoa-Schioppa, C., Pasternak, T., Seo, H., Lee, D., et al. (2014). A hierarchy of intrinsic timescales across primate cortex. Nature Neuroscience 17, 1661–1663. https://doi.org/10.1038/nn.3862

Nambu, A., and Llinaś, R. (1994). Electrophysiology of globus pallidus neurons in vitro. Journal of neurophysiology 72, 1127–1139. DOI: 10.1152/jn.1994.72.3.1127

Nishida, S., Tanaka, T., Shibata, T., Ikeda, K., Aso, T., and Ogawa, T. (2014). Discharge-rate persistence of baseline activity during fixation reflects maintenance of memory-period activity in the macaque posterior parietal cortex. Cerebral Cortex 24, 1671–1685. DOI: 10.1093/cercor/bht031

Nougaret, S., and Ravel, S. (2015). Modulation of tonically active neurons of the monkey striatum by events carrying different force and reward information. Journal of Neuroscience 35, 15214 – 15226. DOI: 10.1523/JNEUROSCI.0039-15.2015

Nougaret, S., and Ravel, S. (2018). Dynamic Encoding of Effort and Reward throughout the Execution of Action by External Globus Pallidus Neurons in Monkeys. Journal of Cognitive Neuroscience 30, 1130–1144. DOI: 10.1162/jocn_a_01277

Nougaret, S., Meffre, J., Duclos, Y., Breysse, E., and Pelloux, Y. (2013). First evidence of a hyperdirect prefrontal pathway in the primate : precise organization for new insights on subthalamic nucleus functions. Frontiers in Computational Neuroscience 7, 1–2. https://doi.org/10.3389/fncom.2013.00135

Ogawa, T., and Komatsu, H. (2010). Differential temporal storage capacity in the baseline activity of neurons in macaque frontal eye field and area V4. Journal of Neurophysiology 103, 2433–2445. DOI: 10.1152/jn.01066.2009

O’Reilly, R.C., and Frank, M.J. (2006). Making working memory work: A computational model of learning in the prefrontal cortex and basal ganglia. Neural Computation 18, 283–328. DOI: 10.1162/089976606775093909

Paladini, C.A., and Tepper, J.M. (2016). Neurophysiology of Substantia Nigra Dopamine Neurons: Modulation by GABA and Glutamate. Handbook of Behavioral Neuroscience 24, 335–359. https://doi.org/10.1016/B978-0-12-802206-1.00017-9

Petroni, F., Panzeri, S., Hilgetag, C.C., Kötter, R., and Young, M.P. (2001). Simultaneity of responses in a hierarchical visual network. NeuroReport 12, 2753–2759. DOI: 10.1097/00001756-200108280-00032

Raz, A., Vaadia E., and Bergman H. (2000). Firing patterns and correlations of spontaneous discharge of pallidal neurons in the normal and the tremulous 1-Methyl-4-Phenyl-1,2,3,6-Tetrahydropyridine vervet model of parkinsonism. Journal of Neuroscience 20, 8559–8571. DOI: 10.1523/JNEUROSCI.20-22-08559.2000

Redgrave, P., Prescott, T.J., and Gurney, K. (1999). The basal ganglia: a vertebrate solution to the selection problem? Neuroscience, 89, 1009–1023. DOI: 10.1016/s0306-4522(98)00319-4

